# Genetics of cortico-cerebellar expansion in anthropoid primates: a comparative approach

**DOI:** 10.1101/043174

**Authors:** Peter W. Harrison, Stephen H. Montgomery

**Author notes:** PWH and SHM contributed equally.

## Abstract

What adaptive changes in brain structure and function underpin the evolution of increased cognitive performance in humans and our close relatives? Identifying the genetic basis of brain evolution has become a major tool in answering this question. Numerous cases of positive selection, altered gene expression or gene duplication have been identified that may contribute to the evolution of the neocortex, which is widely assumed to play a predominant role in cognitive evolution. However, the neocortex co-evolves with other, functionally inter-dependent, regions of the brain, most notably the cerebellum. The cerebellum is linked to a range of cognitive tasks and expanded rapidly during hominoid evolution, independently of neocortex size. Here we demonstrate that, across primates, genes with known roles in cerebellum development are just as likely to be targeted by selection as genes linked to cortical development. In fact, cerebellum genes are more likely to have evolved adaptively during hominoid evolution, consistent with phenotypic data suggesting an accelerated rate of cerebellar expansion in apes. Finally, we present evidence that selection targeted genes with specific effects on either the neocortex or cerebellum, not both. This suggests cortico-cerebellar co-evolution is maintained by selection acting on independent developmental programs.

## Introduction

The proximate basis of primate brain expansion and increased cognitive performance lies in changes in gene function and regulation. Identifying the genetic basis of phenotypic change can provide insights into how developmental mechanisms evolve, how they are constrained, and how changes at a cellular level contribute to broad scale anatomical evolution (1). This potential for dissecting the biological basis of brain and behavioural evolution motivates many genomic comparisons across primates (2). These have identified numerous genes associated with brain development with high rates of evolution (3–9), divergent expression profiles (10–13) or duplicated sequence (14–17), either across primates or during recent human evolution. In several of these cases the genetic changes have been demonstrated to have functional effects on neuronal proliferation or maturation (7,8,15,18,19). These results highlight potential cellular adaptations driving changes in brain size, and provide a powerful means of investigate human-specific adaptations.

The majority of these examples investigate genes linked to neocortical evolution, reflecting the widely held assumption that the neocortex has a predominant role in ‘higher’ cognition (20). However, the neocortex co-evolves with other brain components with which it is functionally connected, suggesting a complete understanding of primate brain expansion will not be found by focusing solely on neocortex development (21–23). Of particular importance is the relationship between the neocortex and cerebellum. Across primates cortico-cerebellar co-evolution pervades biological levels, occurring at a coarse volumetric scale, at the level of individual nuclei, and at the cellular level (21,22,24).

In humans, the cerebellum is increasingly recognised to be important for both motor and ‘higher cognitive’ function, including the capacity to plan and execute complex behavioural sequences (20,23,25). Similarly, across primates, cerebellar expansion is linked to extractive foraging, independently of neocortex volume (23). The importance of the cerebellum in primate brain evolution is further bolstered by comparative analyses that demonstrate a rapid, non-allometric expansion of the cerebellum relative to neocortex size in hominoids that is indicative an adaptive change in the cortico-cerebellar functional relationship (26–29). Hominin evolution is also characterised by reciprocal expansion of the neocotex and cerebellum, with recent modern humans being distinguished from our early modern humans by an increase in cerebellum volume relative to the neocortex (30).

The importance of coordinated cortico-cerebellum expansion suggests that we must look beyond neocortical evolution to obtain a full picture of the genetic architecture of primate brain evolution. The likely action of positive selection on genes involved in cerebellum development is further suggested by the accelerated rate of evolution of *AHI1*, a gene associated with developmental disorders of the cerebellum, during human evolution (31). In addition, genetic approaches may be useful in addressing long-standing debates about the constraints governing co-evolving brain networks. For example, do the neocortex and cerebellum co-evolve through genetically independent developmental changes maintained by selection (*sensu* Barton and Harvey (21))? Or are they the result of pleiotropic genetic effects shared across anatomical boundaries (*sensu* Finlay and Darlington (32))? If the latter is true, do these pleiotropic effects reflect an evolutionary constraint, or have they evolved to maintain the functional relationship between components during brain expansion? These questions will be central to providing a full understanding of brain evolution, and also address questions of long standing interest in evolutionary biology on the trade-offs between adaptation and constraint in the genetic basis of composite traits.

Here we ask three questions that provide an initial assessment of the role of genes controlling the development of the neocortex and cerebellum in anthropoid brain evolution. First, we ask whether genes with known roles in the development of the cerebral cortex, which is predominantly composed of the neocortex, are more likely to be targets of positive selection than those affecting cerebellum development. Second, whether patterns of molecular evolution mirror non-allometric changes in component size. Finally, whether these genes evolve in a manner suggesting a specific evolutionary association with either brain component to explore whether cortico-cerebellar co-evolution is maintained by selection acting on a common or independent set of genes.

## Methods

### Data collection

We obtained a list of human genes with known roles in ‘cerebral cortex development’ or ‘cerebellum development’ from two ontology databases; Amigo (33) (GO:0021987 and GO:0021549) and EBI’s QuickGO (34) (GO:0021987 and GO:0021549). These GO terms have the best-matched definitions among those relevant for each component. In anthropoids, the neocortex comprises the vast majority of the cerebral cortex (~90%; Stephan et al. 1981). This GO search resulted in 198 Amigo and 300 QuickGO genes associated with cerebral cortex development, and 144 Amigo and 222 QuickGO genes associated with cerebellum development, which are not mutually exclusive, that were then combined to form the starting gene set. This starting human gene set was then used to obtain 1:1 orthologs from 11 anthropoid genomes (Figure 1A) using a reciprocal best hit BLASTn (36) approach between each species and the human coding sequence with an e-value cut-off of 1e^-10^ and a minimum percentage identify of 30. 11-way 1:1 ortholog sets were aligned using codon aware PRANK v 140615 (37), converted to phylip format in preparation for PAML analyses (38) and filtered using a conservative alignment filtering program SWAMP v 1.0 (39) to remove regions of poorly aligned or error-rich sequences on a branch-specific basis. SWAMP was run twice, first using a threshold of 5 and a window size of 15, and then a second run with a threshold of 2 with a window size of 3, with a minimum sequence length of 300 bases and *interscan* masking for both runs. The final, strict and conservatively filtered 11-species dataset consisted of three non-overlapping groups: 53 genes with known roles cerebral cortex development, 47 genes with known roles in cerebellum development, and 10 genes with known roles in both (Table S1).

**Figure 1.**
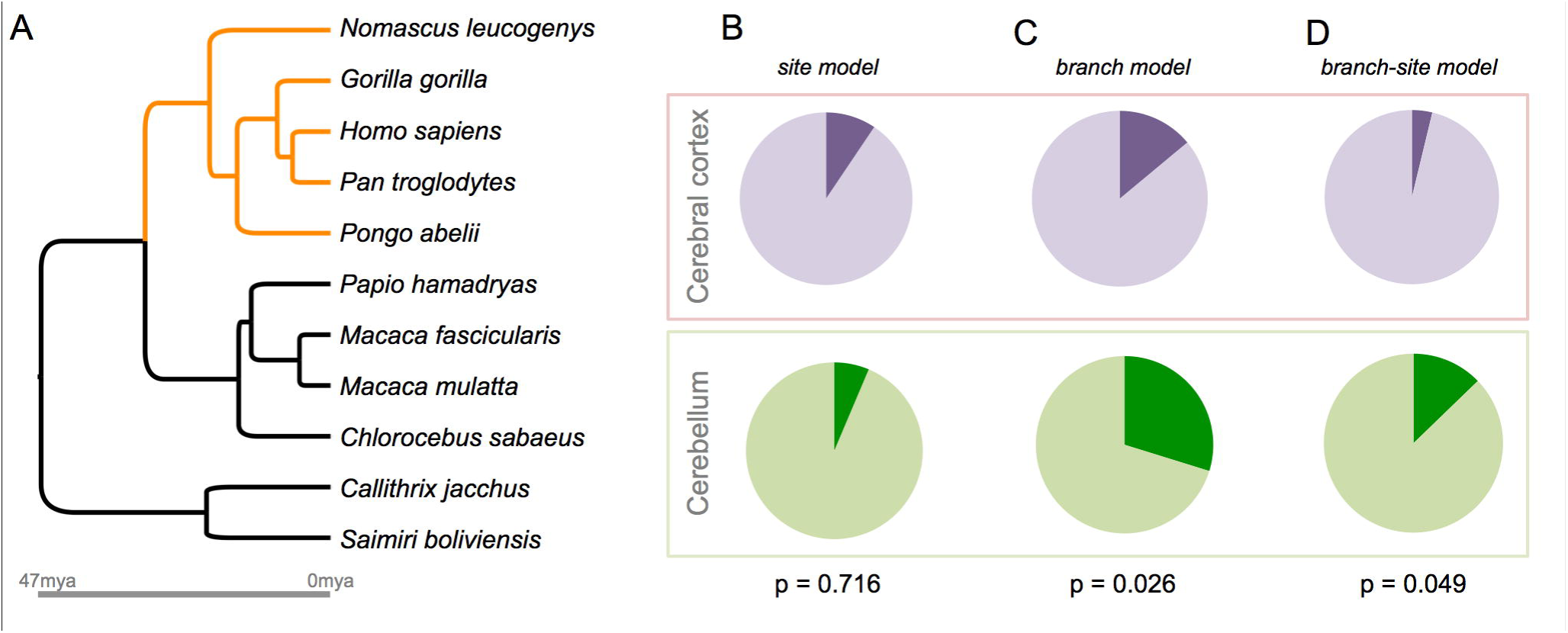
**A)** Phylogeny of the 11 anthropoids included in the study: hominoid lineages are shown in orange. **B-D)** Pie-charts showing the proportion of genes significant (darker colour) under each test for cerebral cortex (red) and cerebellum genes (green). The p-value from a Fisher’s Exact test comparing the groups is shown below.

### Selection analyses

Estimation of *dN/dS* ratios (ω), a common measure of the strength of selection acting on a protein coding gene, was carried out using a codon-based maximum likelihood method (PAML v.4) (38). Nested models were compared using the likelihood ratio test statistic (–2[loglikelihood1 – loglikelihood2]) to critical values of the χ^2^ distribution and degrees of freedom as the difference in the number of parameters estimated in each model. We compared the frequency of positive selection acting on genes associated with cerebral cortex and cerebellum development in two ways. First, we used the site model tests of positive selection (‘Model 8/8a’) to identify genes evolving under positive selection across anthropoids. The site models allow ω to vary across sites, but not across branches. Second, we used the branch-site models (‘new model A’) to identify genes under increased positive selection in hominoids (Figure 1A). The branch-site models allow ω to vary across both sites and branch categories defined a priori. We repeated this test for accelerated rates of evolution in hominoids using a branch model test, where ω is fixed across sites but varies between branch categories. For each analysis, the percentage of each category of gene that experienced selection were compared using a Z-test and more conservative Fisher’s Exact Test.

### Gene-phenotype co-evolution

We sought to test the link between the molecular evolution of our gene-set and the evolution of neocortex and cerebellum size using a phylogenetic, comparative approach (40). Branch models in PAML were used to calculate the root-to-tip *dN/dS* for each species. These were then regressed against the phenotypic trait of interest using a Phylogenetic Generalised Least Squares (PGLS) regression implemented in BayesTraits (41), that corrects for the nonindependence of inter-specific data. Each gene was regressed against neocortex volume and cerebellum volume as predictor variables simultaneously. This permits the identification of genes with either a specific co-evolutionary association with either neocortex or cerebellum size, or genes that co-vary with both traits independently. We also repeated the analyses including rest-of-brain volume as an additional predictor variable, which lead to similar results. In all cases root-to-tip *dN/dS* and brain component volumes were log_10_-transformed.

## Results

### What proportion of genes with known roles in cerebral cortex or cerebellum development are targeted by positive selection?

Across anthropoids, the average rate of evolution of genes with known roles in the development of the cerebellum does not differ from those that function in the development of the cerebral cortex, which is predominantly comprised of neocortex (Mann-Whitney U(_98_) = 1167.5, Z = 0.539, p = 0.590) (Table S2). Genes linked to the development of both brain components do not differ from those specifically linked to cerebral cortex (U(_61_) = 296.5, Z = 0.592, p = 0.554) or cerebellum development (U(_55_) = 241.5, Z = 0.136, p = 0.892). Site-model tests for positive selection acting at a subset of codons also identify a similar proportion of genes associated with cerebral cortex or cerebellum development with evidence of positive selection across anthropoids. 3/47 cerebellum genes (6.4%) are significant at a nominal α of 0.05, compared to 5/53 cerebral cortex genes (9.4%). These proportions are not significantly different (Fisher’s Exact Test, p = 0.716, Z-test: z = 0.5613. p = 0.575) (Table, 1; Figure 1B; Table S3). The same conclusion is reached after correcting for multiple testing using the false discovery rate (FDR) (42), after which the site-model test is significant for two genes linked to cerebellum development (*RPGRIP1L*, *PCNT*) and one gene linked to cerebral cortex development (*TACC2*). None of the 10 genes with annotated function in the development of both brain components show evidence of positive selection. These results suggest genes affecting cerebellum development are just as likely to be targeted by positive selection as those affecting cerebral cortex development.

### Do rates of molecular evolution reflect rates of brain component expansion?

A significantly greater proportion of cerebellum genes (14/47, 29.8%) than cerebral cortex genes (6/53, 11.3%) experienced an accelerated rate of evolution in hominoids, taking a nominal α of 0.05 (Fisher’s Exact Test, p = 0.026, Z-test: z = 2.304, p = 0.021) (Table 1; Figure 1C; Table S4). This result is also reflected in the branch-site test where the proportion of cerebellum genes (6/47, 12.8%) with evidence of episodic positive selection in hominoids again exceeds the proportion of cerebral cortex genes (2/53, 3.8%). In this case the trend does not reach significance (Fisher’s Exact Test, p = 0.143, Z-test: z = 1.654, p = 0.099) (Table S5). However, one assumption of the branch-site test is the absence of positive selection in the background (non-hominoid) branches (43). After excluding genes with evidence of positive selection across anthropoids under the site-model test (Table S3), the trend observed in the branch-site test becomes significant at p < 0.05 (Fisher’s Exact Test, p = 0.049, Z-test: z = 2.136, p = 0.032) (Table 1; Figure 1D). These contrasting proportions suggest the strength of positive selection acting on genes controlling cerebellum development increased during hominoid evolution, a clade in which the rate of cerebellar expansion significantly accelerated (29), without a corresponding acceleration in cerebral cortex expansion (21,44).

**Table 1:**
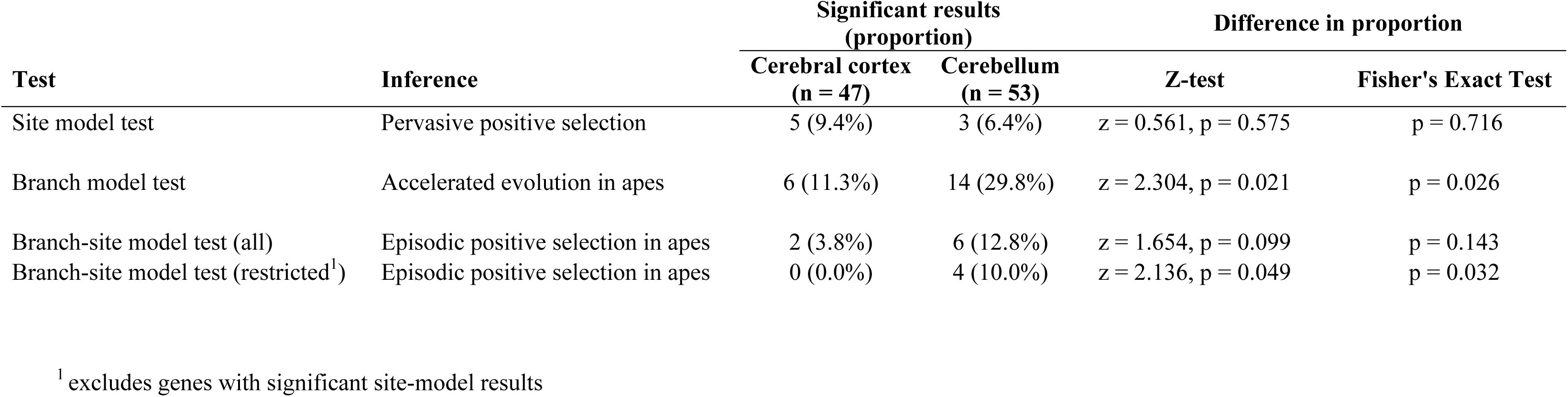
summary of results of *dN/dS* analyses and tests of positive selection.

### Is selection associated with interspecific variation in the size of specific brain components?

We identify 11/47 cerebellum genes (23.4%) that co-evolve with cerebellum volume, after controlling for neocortex volume, of which 4 are significant at p <0.001 and 2 (*RPGRIP1L, ATRN*) remain significant after FDR-correction (Table 1; Figure 2; Table S6). We re-nalaysed the top four genes associated with cerebellum volume, separating *dN* and *dS* whilst controlling variation in neocortex volume, to test if the association is driven by variation in *dN*. For 3/4 genes we find a significant partial regression with *dN* (*ATRN* t_5_ = 3.789, p = 0.006; *EZH2* t_5_ = 3.990, p = 0.005; *KNDC1* t_5_ = 3.586, p = 0.008), the remaining locus showed a non-significant trend (*RGRIP1L* t_5_ = 1.582, p = 0.087). Only one cerebellum gene (2.1%) shows an association with neocortex volume, which is a significantly lower proportion (Fisher’s Exact Test, p = 0.004, Z-test: z = 3.091, p = 0.002). 1/53 cerebral cortex genes (*DICER1*) shows evidence of co-evolution with neocortex volume (1.9%), whilst 6 (11.3%) show evidence of an association with cerebellum size (Table S6). This is not a significant difference in proportion (Fisher’s Exact Test, p = 0.113, Z-test: z = 1.955, p = 0.050), only one of these associations is significant at p < 0.001 and none survive FDR-correction. One of the ten genes annotated as functioning in both neocortex and cerebellum development (*GART*) shows an association with cerebellum, but not neocortex, volume (Table S6). Similar results were obtained when rest-of-brain was included in the regression model. No gene shows an association with variation in both neocortex and cerebellum volume, regardless of the gene category or regression model. These results suggest selection acts on genes associated with phenotypic variation specific to each brain component.

**Figure 2.**
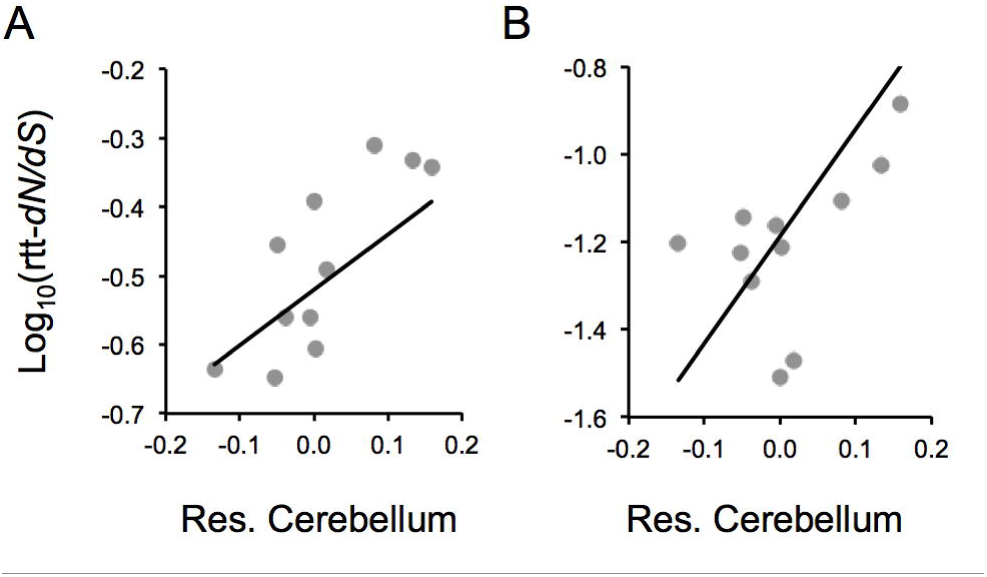
Phenotypic associations for two genes A) *RGRIP1L* and B) *ATRN*. The log_10_-transformed root-to-tip (rtt-) *dN/dS* is plotted against residual cerebellum volume, calculated from a PGLS regression of cerebellum against neocortex volume. The black line shows the result of a phylogenetically-controlled regression between rtt-*dN/dS* and residual cerebellum volume. This figure is for illustrative purposes, in the main analyses the two brain components were included as variables in a multiple regression.

## Discussion

Primate brain expansion reflects increases in the volume and neuron number of co-evolving structures (21). This pattern of distributed adaptation must be reflected in the molecular evolution of genes controlling brain development. Our results provide two contributions to understanding the genetic basis of brain evolution. First, we provide evidence that genes associated with cerebral cortex development are no more likely to have evolved adaptively across anthropoids than cerebellum genes. Indeed, significantly more cerebellum genes experienced an increased rate of evolution in hominoids, consistent with the non-allometric expansion of the cerebellum, but not the neocortex, in apes (29). Second, we provide evidence that the selection regimes shaping these genes are linked to the evolution of specific brain components.

Our results raise the possibility that a significant proportion of the genetic changes that underpin adaptive evolution of primate brain size, structure and cognition will affect aspects of non-cortical development. This conclusion is consistent with evidence of adaptive expansion in cerebellum volume during hominoid evolution (29), and is further supported by a phylogenetic analysis of lineage-specific shifts in gene expression across mammals that found an over-abundance of hominoid-specific expression shifts in genes expressed in the cerebellum (10). These results suggest future comparative studies of primate gene expression should not be limited to samples derived from cortical tissue, and functional tests of candidate genes targeted by positive selection should consider the phenotypic relevance of changes in gene function in non-cortical structures.

Of course, our results are wholly dependent on the quality of gene ontology annotation. Our incomplete knowledge of gene function means that it is inevitable that the genes included in this study reflect a minority of those that influence cerebral cortex and cerebellum development. However, gene ontology datasets are routinely used in post-hoc tests for functional enrichment where the same caveats apply, here we have simply used the available data to facilitate tests of specific hypotheses. We know of no reason why either gene set may be biased in a way that would produce contrasting patterns of results, but cannot formally rule out this possibility. Finally, in this study we adopted a conservative approach, analysing only genes with strict 11-way 1:1 orthologues in published anthropoid genomes and removing short and/or poorly aligned sequence. This further reduces the gene set, but produces more reliable estimates of selection regimes (39).

Our analyses highlight several genes with patterns of molecular evolution that link them to inter-specific differences in cerebellum volume. These include two genes, *RGRIP1L* and *ATRN*, with a particularly strong signal of co-evolution between the strength of selection acting on their coding sequence and cerebellum volume, independently of variation in neocortex or rest-of-brain volume, in our gene-phenotype association tests (Figure 2). *RGRIP1L* is one of a small number of loci linked to Joubert Syndrome, a rare genetic disorder associated with severe hypoplasia of the mid-hindbrain and cerebellum (45,46). The cellular role of *RGRIP1L* appears to be in the correct function of the cilia and basal bodies (47,48) which are necessary for expansion of the cerebellar neural progenitor pool (49). Disruption of *ATRN*, an E3 ubiquitin ligase, in *Mus* causes vacuolization and degeneration of the cerebellum (50,51). Another ubiquitin ligase, *UBE3A*, has been implicated in brain and cognitive development (52) and interacts with *ASPM*, a key regulator of brain size (53). A third gene, *EZH2*, also shows evidence of a phenotypic association with cerebellum volume but is narrowly non-significant after FDR correction. *EZH2*, functions to regulate neuronal migration of precerebellar neurons during the development of the cortico-cerebellar connectivity (54).

We also find multiple genes with significantly accelerated rates of evolution in hominoids, coincident with an accelerated rate of cerebellar expansion (29). Several of these genes also show an association with cerebellum volume in our gene-phenotype association tests at a nominal α threshold of 0.05. These include *AGTPBP1*, disruption of which causes cerebellar Purkinje cell degeneration (55), *PCNT*, which causes primordial dwarfism with microcephaly (56), *TH*, a gene linked to two disorders which affect motor control, Segawa Syndrome and Parkinson’s (57,58), and *MYO16* and *GART* which have putatively been linked to Autistic Spectrum Disorders and Down Syndrome respectively (59,60).

Finally, we identify patterns of molecular evolution that may implicate a separate group of genes in the evolution of neocortex size. Only one locus shows a phenotypic association with neocortex volume; *DICER1*, which regulates neurogenesis in the developing cerebral cortex in a time dependent manner (61,62). Notably, the two genes with the strongest evidence for episodic positive selection in hominoids belong to the same gene family. *TACC1* and *TACC2* are both associated with regulation of nuclear migration and are required for normal patterns of self-renewal in neural progenitors. TACCs interact with a centrosomal protein, CEP120, to regulate nuclear migration and the self-renewal of cortical neural progenitor cells by controlling microtubule growth (63). A similar function is thought to mediate the influence of microcephaly genes on brain development (18,64).

Beyond functional affects of individual genes, our approach has the potential to tackle fundamental questions about how composite or modular tissues, such as the brain, evolve. For example, two models of brain evolution dominate debates surrounding the adaptive significance of variation in brain structure. One model proposes a conserved developmental program drives a ‘concerted’ pattern of brain evolution, with selection shaping the overall size of the system but not individual components(32,65). An alternative model instead argues that different brain regions evolve independently of overall brain size to meet species-specific behavioural needs, resulting in a ‘mosaic’ pattern of brain evolution, but may co-evolve due to functional interdependence (21,66). These two models implicitly make contrasting predictions about the genetic architecture of brain structure. The concerted model predicts variation in the size of different brain regions will be determined by genetic correlations between those structures, i.e. variation at a common set of genes. The mosaic model instead predicts that the development of different brain regions must be at least partially distinct in order to facilitate independent evolution.

In recent years these predictions have been tested using quantitative genetics within a range of vertebrates, either using wild pedigrees (67–69), inbred strains (70) or divergent populations/domestic breeds (71,72). In support of the mosaic brain hypothesis, these have found little evidence for widespread genetic co-variation between major brain components. Similarly, large genome-wide association studies within humans that have identified independent genetic bases associated with brain regions (73,74). Quantitative genetics assesses the phenotypic associations of standing genetic variation within populations and their relevance to macroevolution depends on the relative frequency at which selection acts on *de novo* mutations that may cause different patterns of genetic correlation. Our results therefore complement these intra-specific studies, providing the first inter-specific test designed to identify genes associated with neocortex and/or cerebellum evolution. These two structures show a consistent pattern of co-evolution across primates (21,22,24), reflecting their functional inter-dependence (23,25). Our analyses did not identify any gene that covaries with both structures, but does identify multiple genes with a specific association with either the cerebellum or neocortex. This suggests the coevolution of these structures is unlikely to be solely due to genetic integration or pleiotropy. If more broadly true, this conclusion bolsters the interpretation of cortico-cerebellar coevolution as indicating adaptive co-evolution, maintained by selection acting on distinct developmental pathways.

Comparative functional analysis of the genes highlighted by our analyses will be necessary to confirm and extend these conclusions. These will also be needed to address key questions beyond functional effects. For example, when disrupted, several of the genes highlighted by our analyses affect the development of multiple organs. For example, both *AGTPBP1* and *PAFAH1B* have known roles in spermatogenesis (75,76), whilst disruption of *PCNT* can affect global somatic growth (56). If these genes do have a specific evolutionary role in cerebellar development, how are these pleiotropic effects avoided? Similar questions have been raised over previous candidate genes (3,18), further emphasising the importance of coupling comparative and functional data.

In summary, we have presented an analyses aimed at providing an initial assessment of the strength of selection targeting genes with development roles in distinct brain regions. Although our understanding of gene ontology is incomplete, we illustrate how this information can be used to test hypotheses in a phylogenetic comparative setting, in addition to post-hoc enrichment analyses. We highlight that there is currently no evidence that selection is limited or biased towards genes affecting cerebral cortex development, and encourage evolutionary geneticists to adopt a cohesive view of brain evolution that encompasses the recognised importance or non-allometric expansion of non-cortical regions, and to tackle the central question of co-evolution and relative genetic independence of brain components. Finally, we further illustrate the potential of hypothesis driven comparative genetics in dissecting the genetic basis of phenotypic evolution. The ever-increasing numbers of sequenced genomes will permit increasingly powerful analyses of the targets of selection and their phenotypic relevance.

## Acknowledgements

We thank Amy Boddy, Robert Barton, Nicholas Mundy, Judith Mank and her lab at UCL for comments on draft manuscripts. SHM is grateful for an Early Career Research Fellowship from the Leverhulme Trust.

## References

1. Rausher MD, Delph LF. When does understanding phenotypic evolution require identification of the underlying genes? Evolution. 2015;69(7):1655–64.

2. Enard W. Comparative genomics of brain size evolution. Front Hum Neurosci. 2014;8:345.

3. Montgomery SH, Capellini I, Venditti C, Barton RA, Mundy NI. Adaptive evolution of four microcephaly genes and the evolution of brain size in anthropoid primates. Mol Biol Evol. 2011;28(1):625–38.

4. Montgomery SH, Mundy NI. Evolution of ASPM is associated with both increases and decreases in brain size in primates. Evolution. 2012;66(3):927–32.

5. Montgomery SH, Mundy NI. Positive selection on NIN, a gene involved in neurogenesis, and primate brain evolution. Genes, Brain Behav. 2012;11(8):903–10.

6. Pollard KS, Salama SR, King B, Kern AD, Dreszer T, Katzman S, et al. Forces shaping the fastest evolving regions in the human genome. PLoS Genet. 2006;2(10):1599–611.

7. Kamm GB, López-Leal R, Lorenzo JR, Franchini LF. A fast-evolving human NPAS3 enhancer gained reporter expression in the developing forebrain of transgenic mice. Philos Trans R Soc Lond B Biol Sci. 2013;368:20130019.

8. Boyd JL, Skove SL, Rouanet JP, Pilaz L-J, Bepler T, Gordân R, et al. Human-chimpanzee differences in a FZD8 enhancer alter cell-cycle dynamics in the developing neocortex. Curr Biol. 2015;772–9. A

9. Enard W, Przeworski M, Fisher SE, Lai CSL, Wiebe V, Kitano T, et al. Molecular evolution of FOXP2, a gene involved in speech and language. Nature. 2002;418:869–72.

10. Brawand D, Soumillon M, Necsulea A, Julien P, Csárdi G, Harrigan P, et al. The evolution of gene expression levels in mammalian organs. Nature. 2011;478:343–8.

11. Enard W, Khaitovich P, Klose J, Zöllner S, Heissig F, Giavalisco P, et al. Intra‐ and interspecific variation in primate gene expression patterns. Science. 2002;296:340–3.

12. Bauernfeind AL, Soderblom EJ, Turner ME, Moseley MA, Ely JJ, Hof PR, et al. Evolutionary divergence of gene and protein expression in the brains of humans and chimpanzees. Genome Biol Evol. 2015;7(8):2276–88.

13. Khaitovich P, Muetzel B, She X, Lachmann M, Hellmann I, Dietzsch J, et al. Regional patterns of gene expression in human and chimpanzee brains. Genome Res. 2004;14:1462–73.

14. Burki F, Kaessmann H. Birth and adaptive evolution of a hominoid gene that supports high neurotransmitter flux. Nat Genet. 2004;36(10):1061–3.

15. Florio M, Albert M, Taverna E, Namba T, Brandl H, Lewitus E, et al. Human-specific gene ARHGAP11B promotes basal progenitor amplification and neocortex expansion. Science. 2015;347(6229):1465–70.

16. Zimmer F, Montgomery SH. Phylogenetic analysis supports a link between DUF1220 domain number and primate brain expansion. Genome Biol Evol. 2015;7(8):2083–8.

17. Keeney J, Dumas L, Sikela J. The case for DUF1220 domain dosage as a primary contributor to anthropoid brain expansion. Front Hum Neurosci. 2014;8:1–11.

18. Pulvers JN, Bryk J, Fish JL, Wilsch-Bräuninger M, Arai Y, Schreier D, et al. Mutations in mouse Aspm (abnormal spindle-like microcephaly associated) cause not only microcephaly but also major defects in the germline. Proc Natl Acad Sci USA. 2010;107(38):16595–600.

19. Enard W, Gehre S, Hammerschmidt K, Hölter SM, Blass T, Somel M, et al. A humanized version of Foxp2 affects cortico-basal ganglia circuits in mice. Cell. 2009;137:961–71.

20. Parvizi J. Corticocentric myopia: old bias in new cognitive sciences. Trends Cogn Sci. 2009;13:354–9.

21. Barton RA, Harvey PH. Mosaic evolution of brain structure in mammals. Nature. 2000;405(6790):1055–8.

22. Whiting BA, Barton RA. The evolution of the cortico-cerebellar complex in primates: Anatomical connections predict patterns of correlated evolution. J Hum Evol. 2003;44:3–10.

23. Barton RA. Embodied cognitive evolution and the cerebellum. Phil Trans Roy Soc B. 2012;2097–107.

24. Herculano-houzel S, Sherwood CC. Coordinated scaling of cortical and cerebellar numbers of neurons. Front Neuroanat. 2010;4:1–8.

25. Ramnani N. The primate cortico-cerebellar system: anatomy and function. Nat Rev Neurosci. 2006;7:511–22.

26. Balsters JH, Cussans E, Diedrichsen J, Phillips KA, Preuss TM, Rilling JK, et al. Evolution of the cerebellar cortex: The selective expansion of prefrontal-projecting cerebellar lobules. Neuroimage. 2010;49(3):2045–52.

27. Macleod CE, Zilles K, Schleicher A, Rilling JK, Gibson KR. Expansion of the neocerebellum in Hominoidea. J Hum Evol. 2003;44:401–29.

28. Rilling J, Insel T. Evolution of the cerebellum in primates: Differences in relative volume among monkeys, apes and humans. Brain Behav Evol. 1998(52):308–14.

29. Barton RA, Venditti C. Rapid evolution of the cerebellum in humans and other Great Apes. Curr Biol. 2014;24(20):2440–4.

30. Weaver AH. Reciprocal evolution of the cerebellum and neocortex in fossil humans. Proc Natl Acad Sci USA. 2005;102(10):3576–80.

31. Ferland RJ, Eyaid W, Collura R V, Tully LD, Hill RS, Alnouri D, et al. Abnormal cerebellar development and axonal decussation due to mutations in AHI1 in Joubert syndrome. Nat Genet. 2004;36(9):1008–13.

32. Finlay B, Darlington R. Linked regularities in the development and evolution of mammalian brains. Science. 1995;268(521):1578–84.

33. Carbon S, Ireland A, Mungall CJ, Shu S, Marshall B, Lewis1 S, et al. AmiGO: online access to ontology and annotation data. Bioinformatics. 2009;25(2):288–9.

34. Binns D, Dimmer E, Huntley R, Barrell D, O’Donovan C, Apweiler R. QuickGO: a web-based tool for Gene Ontology searching. Bioinformatics. 2009;25(22):3045–6.

35. Stephan H, Frahm H, Baron G. New and revised data on volumes of brain structures in Insectivores and Primates. Folia Primatol. 1981;35(1):1–29.

36. Altschup SF, Gish W, Miller W, Myers E, Lipman D. Basic Local Alignment Search Tool. J Mol Biol. 1990;(215):403–10.

37. Löytynoja A, Goldman N. webPRANK: a phylogeny-aware multiple sequence aligner with interactive alignment browser. BMC Bioinform. 2010;579.

38. Yang Z. PAML 4: Phylogenetic analysis by maximum likelihood. Mol Biol Evol. 2007;24(8):1586–91.

39. Harrison PW, Jordan GE, Montgomery SH. SWAMP : Sliding Window Alignment Masker for PAML. Evol Bioinform Online. 2014;197–204.

40. Montgomery SH, Capellini I, Venditti C, Barton RA, Mundy NI. Adaptive evolution of four microcephaly genes and the evolution of brain size in anthropoid primates. Mol Biol Evol. 2011;28(1):625–38.

41. Pagel M. Inferring the historical patterns of biological evolution. Nature. 1999;401(6756):877–84.

42. Benjamini Y, Hochberg Y. Controlling the false discovery rate: a practical and powerful approach to multiple yesting. J Stat Soc B. 1995;57(1):289–300.

43. Anisimova M, Yang Z. Multiple hypothesis testing to detect lineages under positive selection that affects only a few sites. Mol Biol Evol. 2007;24(5):1219–28.

44. Barton RA, Venditti C. Human frontal lobes are not relatively large. Proc Natl Acad Sci USA. 2013;110(22):9001–6.

45. Joubert M, Eisenring J, Robb JP, Andermann F. Familial agenesis of the cerebellar vermis A syndrome of episodic hyperpnea, abnormal eye movements, ataxia, and retardation. Neurology. 1969; 19(9):813.

46. Doherty D. Joubert Syndrome: Insights into brain. Semin Pediatr Neurol. 2009;16(3):143–54.

47. Arts HH, Doherty D, Beersum SEC Van, Parisi MA, Letteboer SJF, Voesenek K, et al. Mutations in the gene encoding the basal body protein RPGRIP1L, a nephrocystin-4 interactor, cause Joubert syndrome. Nat Genet. 2007;39(7):882–8.

48. Delous M, Baala L, Saloman R, Laclef C, Vierkotten J, Golzio C, et al. The ciliary gene RPGRIP1L is mutated in cerebello-oculo-renal syndrome (Joubert syndrome type B) and Meckel syndrome. Nat Genet. 2007;39(7):875–81.

49. Spassky N, Han Y, Aguilar A, Strehl L, Besse L, Laclef C, et al. Primary cilia are required for cerebellar development and Shh-dependent expansion of progenitor pool. Dev Biol. 2008;317:246–59.

50. Bronson RT, Donahue LR, Samples R, Kim JH, Naggert JK. Mice with mutations in the mahogany gene Atrn have cerebral spongiform changes. J Neuropathol Exp Neurol. 2001;60(7):724–30.

51. He L, Lu X, Jolly AF, Eldridge AG, Watson SJ, Jackson PK, et al. Spongiform degeneration in mahoganoid mutant mice. Science. 2003;299:710–2.

52. Wilkinson LS, Davies W, Isles AR. Genomic imprinting effects on brain development and function. Nat Rev Neurosci. 2007;8(11):832–43.

53. Singhmar P, Kumar A. Angelman syndrome protein UBE3A interacts with primary microcephaly protein ASPM, localizes to centrosomes and regulates chromosome segregation. PLoS One. 2011;6(5):e20397.

54. Meglio T Di, Kratochwil CF, Vilain N, Loche A, Vitobello A, Yonehara K, et al. Ezh2 orchestrates topographic migration and connectivity of mouse precerebellar neurons. Science. 2013;339(6116):204–7.

55. Lalonde R, Strazielle C, Inserm U, Electronique SDM, Médecine F De. Spontaneous and induced mouse mutations with cerebellar dysfunctions: Behavior and neurochemistry. Brain Res. 2006;1140:51–74.

56. Rauch A, Thiel CT, Schindler D, Wick U, Crow YJ, Ekici AB, et al. Mutations in the Pericentrin (PCNT) gene cause primordial dwarfism. Science. 2008;319:816–9.

57. Haavik J, Toska K. Tyrosine Hydroxylase and Parkinson’ s disease. Mol Neurobiol.1998;16(3):285–309.

58. Ludecke B, Dworniczak B, Bartolome K. A point mutation in the tyrosine hydroxylase gene associated with Segawa’ s syndrome. Hum Genet. 1995;6:123–5.

59. Liu YF, Sowell SM, Luo Y, Chaubey A, Cameron RS, Kim H, et al. Autism and intellectual disability-associated KIRREL3 interacts with neuronal proteins MAP1B and MYO16 with potential roles in neurodevelopment. PLoS One. 2015;10(4):e0123106.

60. Brodsky G, Barnes T, Bleskan J, Becker L, Cox M, Patterson D. The human GARS-AIRS-GART gene encodes two proteins which are differentially expressed during human brain development and temporally overexpressed in cerebellum of individuals with Down syndrome. Hum Mol Genet. 1997;6(12):2043–50.

61. Kawase-koga Y, Otaegi G, Sun T. Different timings of dicer deletion affect neurogenesis and gliogenesis in the developing mouse central nervous system. Deve Dyn. 2009;238(11):2800–12.

62. Davis TH, Cuellar TL, Koch SM, Barker AJ, Harfe BD, McManus MT, et al. onditional loss of Dicer disrupts cellular and tissue morphogenesis in the cortex and hippocampus. J Neurosci. 2008;17:4322–30.

63. Xie Z, Moy LY, Sanada K, Zhou Y, Buchman JJ, Tsai L. Cep120 and TACCs control interkinetic nuclear migration and the neural progenitor pool. Neuron. 2007;56(1)79–93.

64. Thornton GK, Woods CG. Primary microcephaly: do all roads lead to Rome? Trends Genet. 2009;25(11):501–10.

65. Finlay BL, Darlington RB, Nicastro N. Developmental structure in brain evolution. Behav Brain Sci. 2001;24:263-78; discussion 278-308.

66. de Winter W, Oxnard CE. Evolutionary radiations and convergences in the structural organization of mammalian brains. Nature. 2001;409(6821):710–4.

67. Fears SC, Melega WP, Service SK, Lee C, Chen K, Tu Z, et al. Identifying heritable brain phenotypes in an extended pedigree of vervet monkeys. J Neurosci. 2009;29(9):2867–75.

68. Rogers J, Kochunov P, Zilles K, Shelledy W, Lancaster J, Thompson P, et al. On the genetic architecture of cortical folding and brain volume in primates. Neuroimage. 2010;53(3): 1103–8.

69. Rogers J, Kochunov P, Lancaster J, Shelledy W, Glahn D, Blangero J, et al. Heritability of brain volume, surface area and shape: An MRI study in an extended pedigree of baboons. Hum Brain Mapp. 2007;28:576–83.

70. Hager R, Lu L, Rosen GD, Williams RW. Genetic architecture supports mosaic brain evolution and independent brain-body size regulation. Nat Commun. 2012;3:1079.

71. Henriksen R, Andersson L, Jensen P, Wright D. From the jungle to the barn: Independent genetic control for increased brain and body size and Mosaic brain evolution in chickens during domestication. In review.

72. Noreikiene K, Herczeg G, Gonda A, Balazs G, Husby A, Merilä J. Quantitative genetic analysis of brain size variation in sticklebacks: support for the mosaic model of brain evolution. Proc Roy Soc B. 2015;282(1810):20151008.

73. Toro R, Poline J-B, Huguet G, Loth E, Frouin V, Banaschewski T, et al. Genomic architecture of human neuroanatomical diversity. Mol Psychiatry. 2015;20(8): 1011–16.

74. Hibar DP, Stein LP, Renteria ME, Arias-Vasquez A, Desrivières S, Jahanshad N, et al. Common genetic variants influence human subcortical brain structures. Nature. 2015;520(7546):224–229.

75. Kim N, Xiao R, Choi H, Jo H, Kim J, Uhm S, et al. Abnormal aperm development in pcd 3J ‐ / ‐ Mice: the Importance of Agtpbp1 in spermatogenesis. Mol Cell. 2011;(2005):39–48.

76. Yan W, Assadi AH, Wynshaw-boris A, Eichele G, Matzuk MM, Clark GD. Previously uncharacterized roles of platelet-activating factor acetylhydrolase 1b complex in mouse spermatogenesis. Proc Natl Acad Sci USA. 2003;100(12):7189–94.

